# Ion Mobility Separation of Isomeric Acyl-lysine Marks in Peptides

**DOI:** 10.64898/2026.01.20.700525

**Authors:** Francis Berthias, Nurgül Bilgin, Dale A. Cooper-Shepherd, Nesrin Hammami, James I. Langridge, Jasmin Mecinović, Ole N. Jensen

## Abstract

Isomeric post-translational modifications (PTMs) on proteins challenge proteomic analyses due to their identical mass and fragmentation patterns. We evaluate high-resolution ion mobility spectrometry (IMS) for separating three naturally occurring acyl-lysine isomer pairs on histones: crotonyl/methacryl, butyryl/isobutyryl and L-/D-lactyl. These PTMs were chemically installed on lysine residues 9 and 18 (K9, K18) of synthetic histone H3 peptides (residues 3-15 and 3-25). Using trapped IMS (TIMS) we observe half-height separation of H3_[3-15]_ peptides possessing crotonyl/methacryl and L/D-lactyl marks, and the lactyl isomers of H3_[3-25]_ can be distinguished. In contrast, multi-pass cyclic IMS (cIM) achieves baseline or near-baseline resolution for every pair, except the longest butyryl/isobutyryl peptide isomers, despite their collision-cross-section differences of ≈1%. We show that resolution increased with the square root of cIM pass number, allowing baseline separation within 300 ms. Beyond separation, structure-mobility relationships emerge: branched modifications (isobutyryl, methacryl) yield more compact gas-phase conformations than their linear analogs (butyryl, crotonyl). For the doubly crotonylated/methacrylated peptides studied, both PTM identity and site determine the mobility. These results demonstrate IMS as a sensitive method for elucidation of acyl-modified histone peptide fine structure by resolving isomeric PTM ambiguity. This addresses a persistent analytical bottleneck and should be included in routine proteomics MS-based workflows

## INTRODUCTION

Post-translational modifications (PTMs) modulate protein activity by affecting conformation, stability, and interaction networks, thus maintaining metabolic pathways and cellular homeostasis.^1^ In histones, PTMs encode regulatory information within chromatin, including the epigenetic code (“histone code”) governing chromatin architecture and gene expression.^2^ Among the many different PTMs on histones, lysine acetylation is a functionally prominent mark involved in protein-protein and protein-DNA interactions, impacting the chromatin structure and function by the neutralization of the positive charge of the ε-amino group on lysine residues.^3^ Beyond acetylation, a broader spectrum of lysine acylation modifications exists on histones,^4^ such as propionylation,^5, 6^ butyrylation,^5^ crotonylation,^7, 8^ methacrylation,^9^ malonylation,^10, 11^ succinylation,^11, 12^ glutarylation^13^ and lactylation,^14^ which have gained interest due to their emerging biological significance in both physiological and pathological contexts.^15–20^ Notably, several of these recently discovered acyl-lysine modifications exist as isomers: D-/L-lactylation (KD-la/KL-la),^14^ linear/branched butyrylation (Kbu/Kibu),^21^ and crotonylation/methacrylation (Kcr/Kmea),^9^ whose individual metabolic origins and regulatory roles remain elusive.^7, 9, 22^

Mass spectrometry (MS) underpins the discovery and mapping of PTMs, exploiting diagnostic mass shifts and fragmentation patterns.^23–25^ Positional isomers of PTMs (same PTM(s) on different residues) can be identified based on characteristic fragment ions in MS^n^ workflows.^26^ However, constitutional, geometric, or stereoisomeric PTMs (isomeric PTMs) installed on the same residue share identical mass and backbone fragments. Thus, routine collision- (CID/HCD) and electron-based (ECD/ETD) fragmentations produce near-identical spectra,^27^ resulting in silent mis-annotations in proteomic datasets.

Targeted strategies exist to resolve individual acyl-lysine isomers, relying on specific reagents or chemical modifications. For example, Ozone-induced dissociation distinguishes crotonyl- from methacryl-lysine by cleaving the α,β-unsaturated bond.^9^ Immuno-affinity enrichment with isomer-specific antibodies and enzymatic assays employing stereospecific delactylases identify L-, D-, and ε-carboxyethyl lactyl marks.^28, 29^ Separation strategies like liquid chromatography (LC) commonly used in proteomics workflows help resolve peptide mixtures. Nevertheless, the subtle polarity and hydrophobicity differences of isomeric PTMs in peptides are difficult or impossible to resolve solely by chromatography, leading to co-eluting PTM peptide isomers. For instance the H4K91cr (residues 80-92) peptide elutes earlier than H4K91mea in standard reversed-phase LC,^9^ while digesting peptides into individual amino acids followed by chiral derivatization with Mosher’s reagent is necessary to separate D- and L-lactyllysine isomers, otherwise inseparable as peptides.^28^

Recent high-resolution ion mobility spectrometry (IMS) platforms such as trapped IMS (TIMS),^30^ cyclic IMS (cIM),^31^ structures for lossless ion manipulations (SLIM),^32^ or field-asymmetric IMS (FAIMS),^33^ have achieved percent/sub-percent collision-cross section (CCS) resolution. In IMS, gas-phase ions travel through an inert gas under the influence of an electric field. Ions are separated based on their ion mobility coefficient (K), which depends on the ions charge, size, shape, and from their interactions with atoms of the drift gas. Under low electric field conditions, K can be converted to rotationally averaged CCS, an instrument-independent structural ion descriptor, suitable for library matching, prediction and calculation.^34–39^ Although deep-learning models can now predict peptide CCS with minor error, their reliability for isomeric PTMs remains untested. Because IMS separation takes place downstream of ionization on millisecond-second timescale, it provides an orthogonal, reagent- and derivation-free dimension to standard MS workflows.^40^

High-resolution operation of TIMS and cIM has been established previously, including the initial demonstrations of the TIMS principle^41, 42^ and multipass cIM architectures capable of extended path lengths.^43^ These instrumental developments underpin the high-resolution separations used here. IMS has resolved a variety of peptide and protein isomers (Figure 1). It includes top-down separation of positional acetyl isomers in histone H4,^44, 45^ characterizations of isomeric histone tails with FAIMS and TIMS–ECD in the middle-down range,^46, 47^ positional phospho-isomers in tau peptides^48^ and in broader phosphopeptide libraries,^49^ and discrimination of D/L- and aspartic/isoaspartic acid isomer in a 11-mer amyloid-β peptide.^50^ We recently reported that TIMS can fully resolve Nπ- and Nτ-methyl-histidine isomers in a 16-mer β-actin fragment peptide,^51^ and baseline resolution of a 72-residues crustacean hyperglycaemic hormone incorporating a D-amino acid substitution.^52^ These examples underscore the capacity of IMS to separate positional, constitutional and stereoisomeric PTMs, but its application to isomeric acyl-lysine PTMs remains largely unexplored. For example, Gómez *et al.* demonstrated that TIMS can separate propionyl-lysine from the chemically distinct acrolein-lysine in a PFGK tetrapeptide.^53^

**Figure 1.**
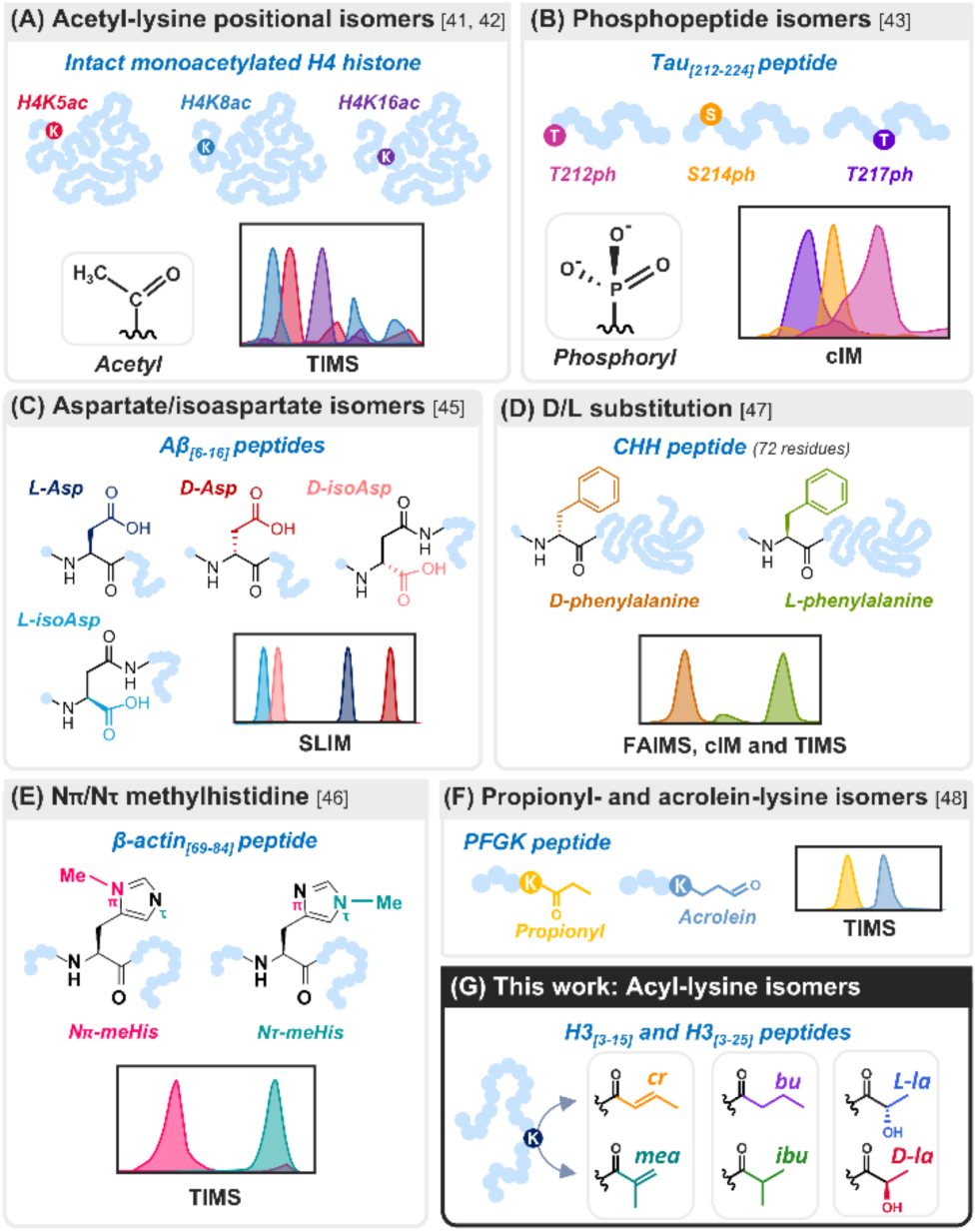
IMS separation of diverse peptide and protein isomers. (A) Acetyl-lysine positional isomers in intact histone H4 proteoforms by TIMS.^44, 45^ (B) Tau phosphopeptide positional isomers (Tau_[212–224]_).^48^ (C) Aspartate/isoaspartate stereoisomers (L-Asp, D-Asp, L-isoAsp, D-isoAsp) in amyloid-β peptide (Aβ_[6–16]_) by SLIM.^50^ (D) D/L substitution in a 72-residue crustacean hyperglycaemic hormone (CHH) by SLIM.^52^ (E) Nπ- and Nτ-methylhistidine in a β-actin peptide (β-actin_[69–84]_) by TIMS.^51^ (F) Propionyl- and acrolein-lysine in a PFGK tetrapeptide by TIMS.^53^ (G) This work: acyl-lysine isomer pairs (crotonyl/methacryl, butyl/isobutyl, L-/D-lactyl) in histone H3 peptides (H3_[3-15]_ and H3_[3-25]_). All panels show schematic structures and IMS peak cartoons.

Here, we address the separation capabilities of two high-resolution IMS platforms: TIMS (timsTOF Pro 2, Bruker Daltonics) and cIM (SELECT SERIES Cyclic IMS, Waters) applied to three pairs of biomedically important isomeric acyl PTMs on lysine residues in H3 peptides: crotonyl/methacryl (Kcr/Kmea; +68.02 Da), butyryl/isobutyryl (Kbu/Kibu; +70.04 Da) and L-lactyl/D-lactyl (KL-la/KD-la; +72.02 Da). Synthetic C-terminally amidated histone peptides H3 (residues 3-15, TKQTARKSTGGKA and residues 3-25, TKQTARKSTGGKAPRKQLATKAA) were prepared with acyl-modifications installed at the naturally existing K9 and/or K18 sites. The Nε-acyl-lysine side-chain structures are shown in Figure 2A (Kcr/Kmea), Figure 3A (Kbu/Kibu), and Figure 4A (KL-la/KD-la). These model peptides allowed us to assess and quantify how unsaturation, branching and chirality of isomeric PTMs affect IMS separation independent of confounding amino acid sequence effects.

**Figure 2.**
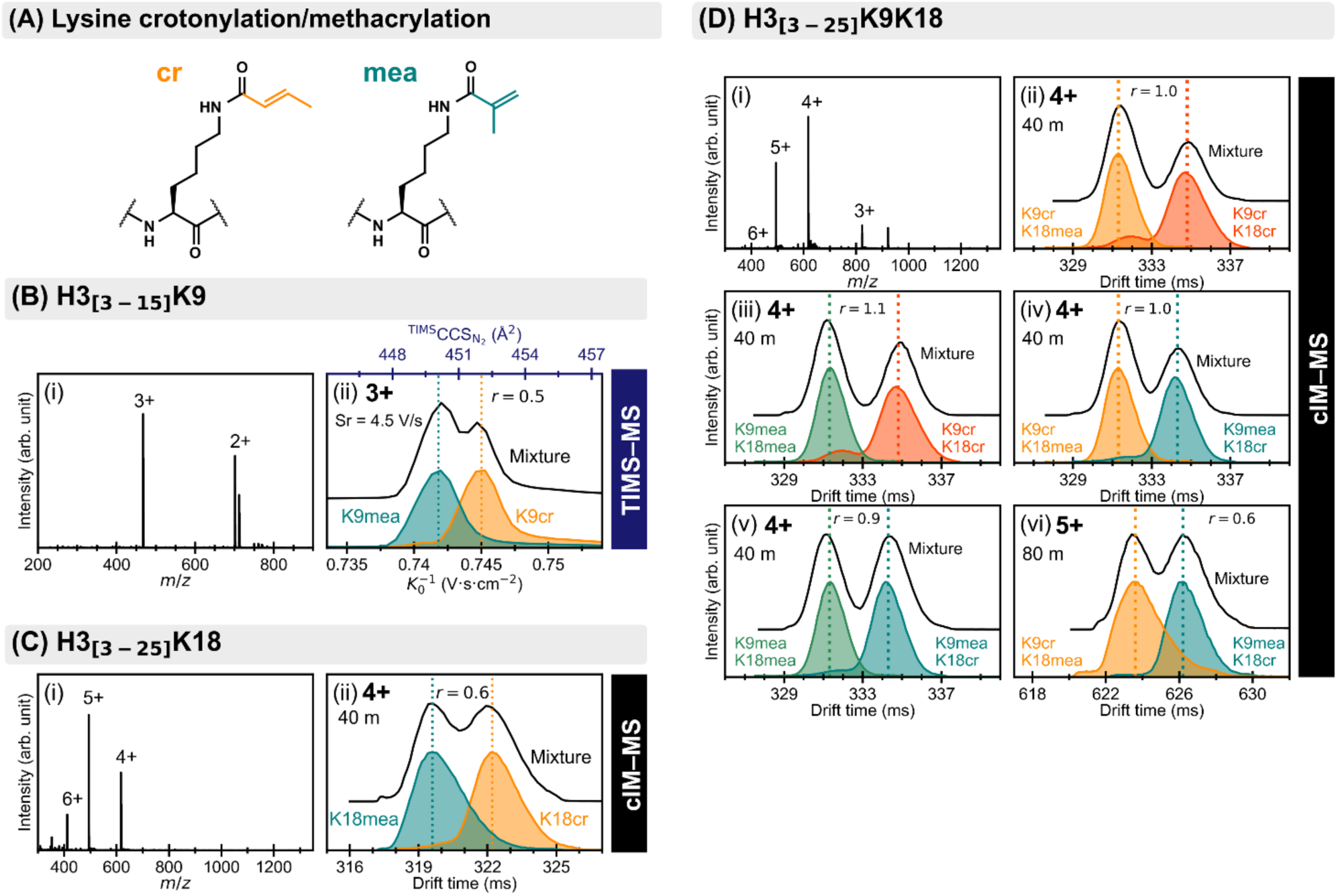
Isomeric crotonyllysine (Kcr) and methacryllysine (Kmea). (A) Nε-acyl-lysine side-chain structures for Kcr (orange) and Kmea (teal). (B) H3_[3-15]_K9cr/K9mea: (i) mass spectrum (timsTOF Pro 2); (ii) TIMS–MS separation of triply charged using a scan rate of 4.5 V/s. (C) H3_[3-25]_K18cr/K18mea: (i) mass spectrum (SELECT SERIES cyclic IMS); (ii) cIM separation after 40 cIM passes for charge state 4+ ions. (D) Doubly modified H3_[3-25]_K9K18: (i) mass spectrum; (ii-v) cIM–MS separations of the isomeric pairs for charge state 4+ after 40 passes: (ii) K9crK18mea vs. K9crK18cr; (iii) K9meaK18mea vs. K9crK18cr; (iv) K9crK18mea vs. K9meaK18cr; (v) K9meaK18mea vs. K9meaK18cr; (vi) Separation of H3_[3-25]_K9crK18mea vs. H3_[3-25]_K9meaK18cr for charge state 5+ after 80 passes. cIM-MS spectra are plotted against drift time (ms); TIMS-MS spectra are plotted as inverse reduced mobility coefficient (K_0_^-1^, lower x-axis) with corresponding CCS values (upper x-axis). Mixtures in black.

**Figure 3.**
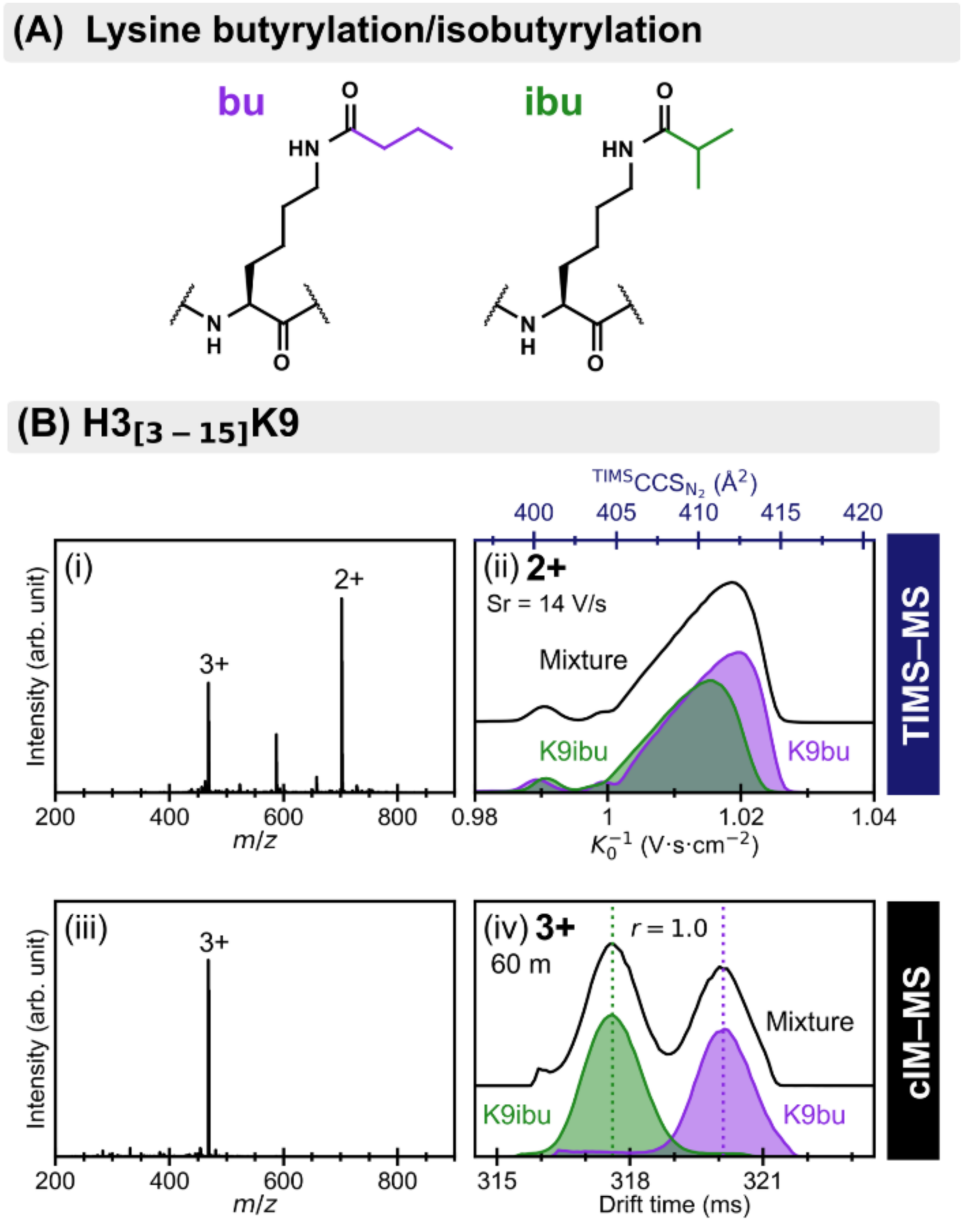
Isomeric butyryllysine (Kbu) and isobutyryllysine (Kibu). (A) Nε-acyl-lysine side-chain structures for Kbu and Kibu. (B) H3_[3-15]_K9bu/K9ibu: (i) mass spectrum (timsTOF Pro 2); (ii) TIMS spectra of doubly charged ions (Sr = 14 V·s^-1^); (iii) mass spectrum (SELECT SERIES cyclic IMS); (iv) cIM baseline separation of triply charged ions after 60 passes. Kbu in violet, Kibu in green, and mixtures in black.

**Figure 4.**
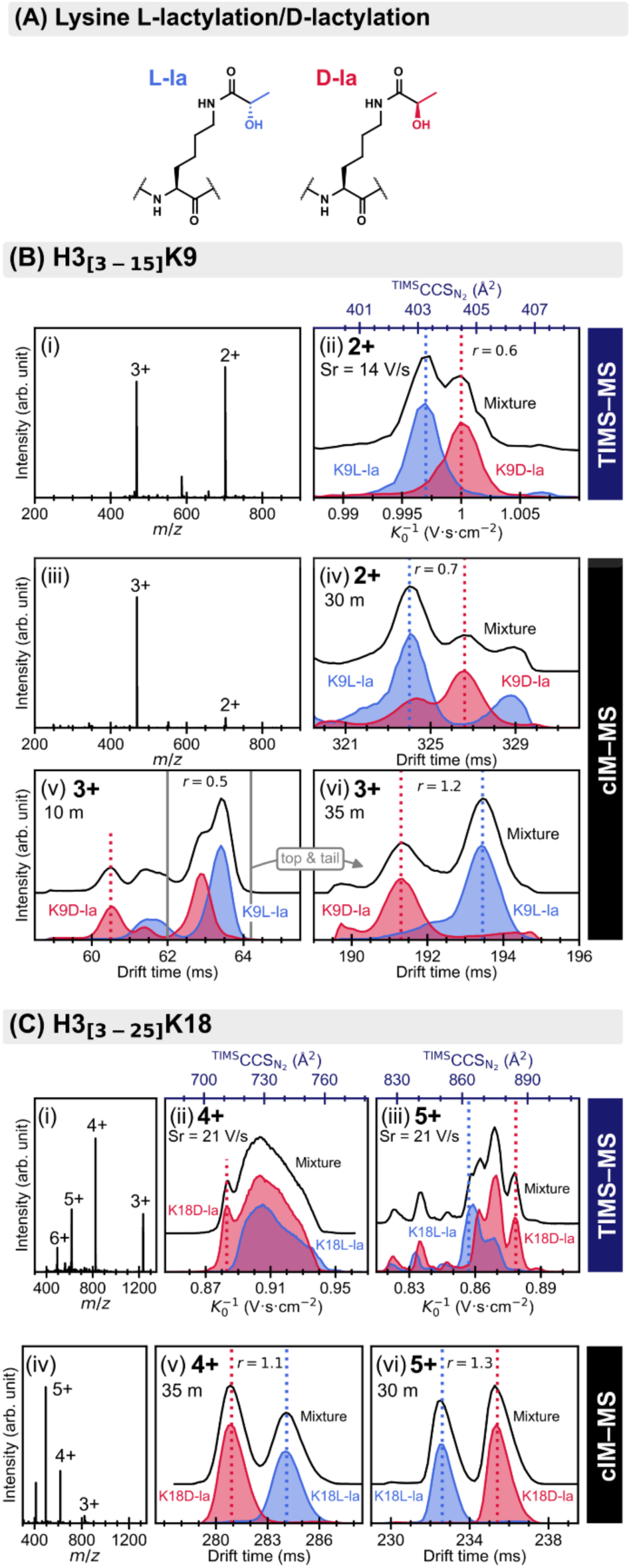
Isomeric L-lactyllysine (KL-la) and D-lactyllysine (KD-la). (A) Nε-acyl-lysine side-chain structures. (B) H3_[3-15]_K9L-la/K9D-la: (i) mass spectrum (timsTOF Pro 2); (ii) TIMS separation of doubly charged ions (Sr = 14 V.s^-1^); (iii) mass spectrum (SELECT SERIES cyclic IMS); (iv)–(vi) cIM separations: 2+ ions after 30 passes, partial separation of 3+ ions (10 m), and their baseline separation after top-and-tail isolation (+25 passes). (C) H3_[3-25]_K18L-la/K18D-la: (i) mass spectrum (timsTOF Pro 2); (ii) and (iii) partial separation of 4+ and 5+ ions (Sr = 21 V·s⁻¹); (iv) mass spectrum (cIM); (v)–(vi) cIM baseline separation of 4+ and 5+ ions after 35 m and 30 m, respectively.

## MATERIALS AND METHODS

### Peptide synthesis

All synthetic histone H3 peptides were prepared on rink amide resin by Fmoc-based solid-phase peptide synthesis (SPPS) employing an orthogonal alloc group on K9 and/or K18 positions. The H3 peptides were selectively acylated using different reagents by performing on-resin reactions on the K9 and/or K18 positions after orthogonal removal of the alloc group. Subsequently, the histone peptides were cleaved from resin, purified by RP-HPLC, and characterized by MALDI-TOF-MS.

### IMS-MS Sample preparation

Purified peptides were dissolved in H₂O/MeOH/HCOOH (49:50:1) at 1-3 µM for direct direct nanoelectrospray ionization (nESI) infusion.

### Trapped ion mobility spectrometry (TIMS)

TIMS-MS experiments were performed on a timsTOF Pro 2 (Bruker Daltonics) with nESI (1.1 kV). Instrumental parameters were optimized for high resolving power as described in previous studies (TIMS in/out tunnel pressure = 3.4/0.84 mbar, ramp time = 1.3 s, accumulation time = 40 ms).^44, 45^ Reduced inverse mobilities and mass-to-charge ratio were calibrated using ESI-L low-concentration tune mix (Agilent), and Collision cross sections values were derived via the Mason–Schamp equation.

### Cyclic ion mobility spectrometry (cIM)

Experiments were carried out on a ESI-Q-cIM-ECD-TOF instrument (Waters) with a drift gas pressure of 1.8 mbar N₂, Travelling wave (TW) velocity of 375 m/s, and TW height of 22 V. Multipass separations were performed to enhance resolution, with resolving power scaling as (number of passes)^1/2^.^31^

### Data analysis

IMS resolving power, Rp, was determined from the ratio of inverse mobility (TIMS) or drift time (cIM) at the peak apex (x_i_) and the corresponding full width at half-maximum (FWHM):

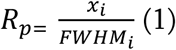

using Compass DataAnalysis (Bruker Daltonics) and MassLynx (Waters). Resolution, r, between isomers was calculated assuming Gaussian peak shapes:

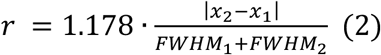

Additional experimental details, including peptide synthesis protocols, calibration procedures, and derivations of IMS resolution equations, are provided in the Supporting Information.

## RESULTS AND DISCUSSION

### Peptide selection for acyl-modified histone analysis

Histone lysine acylations are most abundantly studied and enriched on the N-terminal tails of H3 and H4 proteins. Multiple lysine residues on H3, including K9, K14, K18, K23, and K27 are frequently modified by acylations, with K9 and K18 being heavily acylated *in vivo* and function as recognition sites for histone reader-domains.^9, 54–56^ For this reason, a set of H3 peptides possessing acyl marks at the naturally occurring K9 and K18 sites was synthesized in pairs (Table S1); H3_[3-15]_K9cr/mea, H3_[3-15]_K9bu/ibu, H3_[3-15]_K9D/L-la, H3_[3-25]_K18cr/mea, H3_[3-25]_K18bu/ibu, H3_[3-25]_K18D-/L-la. Pairs of doubly modified H3_[3-25]_K9K18 peptides were also prepared in all four cr/mea isomer permutations (K9cr/K18cr, K9mea/K18mea, K9cr/K18mea and K9mea/K18cr). The shorter peptide (residues 3-15) matches the minimal ARKS lysine-centered sequence used in acyltransferase assays,^57, 58^ whereas the longer peptide represents a longer peptide including the second regulatory lysine residue. All histone peptides were synthesized using standard Fmoc-based solid-phase peptide synthesis (SPPS), with acyl groups site-specifically introduced at the selected lysine residues on-resin after an orthogonal removal of the alloc-protecting group on lysines.

### Charge-state distributions across IMS–MS platforms

nESI-MS analysis of modified peptides was performed using the timsTOF Pro 2 and the SELECT SERIES cyclic IMS instruments. Both instruments generated doubly and triply charged ions for H3_[3-15]_ peptides (mass spectra in Figures 2B-i, 3B-i. 3B-iii, 4B-i and 4B-iii) and charge states ranging from 2+ to 6+ for H3_[3-25]_ peptides (Figure 2C-i, 4C-i and 4B-iv). All mass spectra shown correspond to acquisitions performed with IMS enabled. For the cIM platform, the spectra reflect the single pass (n = 1) condition. The nanolockspray nanoESI source of the SELECT SERIES cIM favours higher charge states; indeed the triply charged ions represent more than 90% of the total ion signal for H3_[3-15]_, whereas the mean charge for H3_[3-25]_ is 4.8 for cr/mea, bu/ibu and L-la/D-la isomers. The nature of the PTMs did not affect the charge state distributions. In contrast, the CaptiveSpray source on the timsTOF Pro 2 produces ∼1:1 ratio of 2+/3+ for H3_[3-15]_ and lower mean charge of 3.2 for H3_[3-25]_ peptide for the same isomeric pairs. These differences likely reflect the higher desolvation capability in the Select Series cIM ESI source.

### Separation of singly and doubly crotonylated/methacrylated histone peptides

Crotonylation and methacrylation differ by the substitution pattern on their α,β-unsaturated carbonyl group (Figure 2A). The crotonyl group features a linear/extended chain with a trans double bond, while the methacryl group possesses a methyl substituent at the α-position, introducing a branch. Figure 2B and 2C presents IMS separation of singly modified cr/mea H3_[3-15]_ and H3_[3-25]_ peptides, respectively. The TIMS spectrum reveals that the triply charged ions exhibit a near half-height separation (peak-to-peak resolution, r ∼ 0.5) with less than 10% signal overlapping at the apex in a 1:1 mixture (Figure 2B-ii) using a scan rate of 4.5 V.s^-1^. The presence and separation of both species is confirmed by the peptide mixture trace (black). For the longer peptide only cIM provides significant separation. After 40m (40 passes) of path length, the two isomers at charge state 4+ can be half-height separated with r = 0.6 (Figure 2C-ii). Higher charge states (5+, 6+) remain unresolved for the same number of passes (Figure S2-vi and vii). TIMS gives no discernible separation for any charge state of H3_[3-25]_ (Sr = 14 V.s^-1^; Figure S2-i-iv), the 4+ distribution spans over 25 Å² which is too broad to reveal any sub-percent ΔCCS between isomers, and the presence of an overlapping multimodal contribution for 5+ ions is also observed.

Resolving doubly modified peptides is analytically valuable as endogenous histone N-terminal tails frequently harbor combinatorial patterns of PTMs, known as PTM-interplay and crosstalk.^59^ We examined all six pairwise combinations of crotonyl- and methacryl-modified H3_[3-25]_K9K18 peptides. TIMS reveals only minor separation for charge state 3+, with K9crK18mea/K9crK18cr, K9meaK18mea/K9crK18ce, and K9meaK18mea/K9crK18cr pairs yielding broadened mixture peaks (Figure S4). No separation is observed for charge states 4+ and 5+ (Figures S5 and S6). Using cIM, the best separation is achieved for charge state 4+, at 40 cIM passes, affording a baseline separation (r∼1) for four of the combinations (Figure 2D-ii-v). The peak order observed in TIMS is consistent with that from cIM. Modification at K18 governs mobility more strongly than modification at K9. Indeed, peptides that differ only at K9 but share the same PTM at K18 partially overlapped (K9crK18cr/K9meaK18cr and K9crK18mea/K9meaK18mea pairs, Figure S3). In contrast, when K9 carried the same PTM, distinct arrival-time distributions emerged depending on whether K18 was modified with crotonyl or methacryl group. The position of the PTM within the peptide sequence modulates its impact on overall conformation.

Between homotypic peptides (K9meaK18mea vs K9crK18cr) the separation is maximal. The branched/branched (K9meaK18mea) peptide drifts 3.3 ms (∼1% of difference) faster than its linear/linear isomeric analog (K9crK18cr), giving a resolution of 0.9-1.1, nearly double the resolution observed for the singly modified K18 pair under identical conditions (Figure 2C-ii). The heterotypic isomers further illustrate the asymmetric contribution of each lysine site, K9meaK18cr elutes 2.9 ms (0.9% difference) earlier than K9crK18mea, whereas fixing K18 as crotonyl reduces the difference between K9meaK18cr and K9crK18cr to 0.6 ms (0.2%) which is insufficient for baseline separation. For the charge state 5+, none of the isomer ions are separated with 40 passes, however, extending the path length up to 80 passes for the pair K9meaK18cr and K9crK18mea lead to a near half-height separation of the two isomers (r ∼ 0.6) (Figure 2D-vi).

### Separation of butyrylated and isobutyrylated histone peptides

Butyryl and isobutyryl groups are structural isomers differing only by an α-methyl branch (Figure 3A). H3_[3-15]_K9bu/ibu ions appear as 2+/3+ (1:0.6) mixture on the timsTOF Pro 2 and predominantly as triply charged on the cIM-MS platform (Figure 3B-i and -iii). Using TIMS, both isomers remain unresolved even at a scan rate of 14 V.s-1 (Figure 3B-ii and Figure S7-i). Sixty cIM passes yield a baseline resolution (r = 1.0) for the 3+ isomers (Figure 3B-iv), with a resolving power of ∼210–220. In contrast, the doubly charged isomers remain unseparated even after extended cIM path length of 80 m (Figure S7-ii). Charge state 4+ (Figure S7-iii) detected at very low intensity for the H3_[3-15]_K9bu and H3_[3-15]_K9ibu peptides yield distinct maxima after 15 passes, but mutual overlap is still present at each apex (∼75.4 ms and ∼77.4 ms).

Regarding the H3_[3-25]_ peptides, neither cIM nor TIMS achieved enough resolution to fully distinguish between K18bu and K18ibu isomers (Figures S8 and S9) despite extended cIM pass number (up to 70 passes for charge state 5+) and optimized TIMS scan condition (21 V.s^-1^). The 4+ distribution spans over 25 Å² which is too broad to reveal any sub-percent ΔCCS between isomers, and the presence of an overlapping multimodal contribution for 5+ ions is also observed. This observation suggests the presence of multiple gas-phase conformations. These pairs of isomers thus appear to be the most challenging to separate in our study.

### Separation of L-/D-lactylated histone peptides

L- and D-lactylations are stereoisomeric acyl modifications that differ only in their stereochemistry of the β-hydroxyl moiety (Figure 4A). The TIMS separation of H3_[3-15]_K9L-/D-la pair, full width at half maximum separation of 2+ ions is achieved with a resolution of 0.6 using a scan rate of 14 V.s^-1^ (Figure 4B-ii). The CCS difference (ΔCCS) is 0.4% between the two doubly charged isomers. To exemplify the separation challenge encounter, this difference is in the same order of magnitude, and even smaller, than the one measured for the PTM positional isomers of intact monoacetylated histone H4 (102 residues, 11.4 kDa).^45^ Using cIM, 30 passes are required to resolve doubly charged H3_[3-15]_K9D-la from H3_[3-15]_K9L-la with a resolution r = 0.7 (Figure 4B-iv). The drift time distribution of H3_[3-15]_K9L-la exhibits overlapping contributions of about 30% of H3K9D-la at its apex, and a secondary peak isolated near 329 ms. For charge state 3+ (Figure 4B-v), 10 passes produce two main peaks partially separated with a resolution of 0.5. To enhance the separation, without racetrack effect of the slowest ions, ions with a drift time between 62-64 ms are selected (“top-and-tail” method) and all the other species are excluded. The selected ions can then be subjected to additional passes; in total 35 passes are necessary to reach a baseline separation (r ∼ 1.2) with a resolving power of ∼185 for both peaks, as shown in Figure 4B-vi.

For the longer lactylated H3_[3-25]_ peptides, TIMS inverse mobility/CCS distributions span a broad envelope of ∼25 Å² and display overlapping features for 4+ and 5+ charge states (Figure 4C-ii and -iii), consistent with the behavior of the H3_[3-25]_K18bu/ibu isomers. Nevertheless, TIMS separation provides resolution of the K18D-la isomer as highlighted with vertical dotted red lines (Figure 4C-ii). K18L-la at charge state 5+ can also be distinguished by a shoulder located at 864 Å² (Figure 4C-iii). cIM achieves a baseline resolution for both 4+ and 5+ ions after 30 and 35 passes with r = 1.1 and 1.3, respectively (Figure 4C-v and -vi). Other charge states exhibit a partial separation (2+ after 10 passes, Figure S10B-iv) or no separation, even after extended acquisition (50 passes for charge state 6+, Figure S10-vi). In general, we observe that the drift time of L- and D-lactyl isomers are reversed for some charge states for both peptide lengths, possibly reflecting different conformational states that likely depend on the protonation sites within ions.

### Influence of carbon-skeleton branching on the ion mobility

The branched isobutyrylated H3_[3-15]_ peptide drifts about 0.8% faster than its linear butyryl isomer in cIM (Figure 3B-iv). A similar trend is observed using TIMS, despite lower resolution, where the butyryl isomer is extended toward slightly higher CCS value (Figure 3B-ii). A similar effect appears for unsaturated acyl groups, indeed the branched methacryl isomer H3_[3-25]_ has a shorter drift time than the linear crotonyl (Figure 2C-iv). In addition, with TIMS, triply charged H3_[3-15]_K9mea shows a lower K0^-1^/CCS value (Figure 2B-ii) and this trend persists for the 2+ ions, although the ΔCCS is too subtle to induce two distinct maxima in the mixture trace (Figure S1). Because CCS represents the orientationally averaged effective surface area of an ion, smaller CCS values for the branched species imply a correspondingly more compact gas-phase conformation. Such branch-induced compaction has been reported for non-peptidic systems such as polyester and PFAS.^60, 61^ For biomolecules, Gomez *et al.* reported a 1% ion mobility difference using TIMS on tetrapeptide isomers bearing propionyl- and acrolein-lysine, differing only by the position of the carbonyl group.^53^ Our results confirm that this steric principle persists even in longer amino acid sequences that remain distinguishable.

### Resolution in multipass cIM

In the cIM analyzer, the drift path length (L) increases with the number of passes (n) (L = Lpass⋅n, where Lpass is the length per pass ≈ 1 m). Two opposing processes influence the resolution for two ions of mobility K_1_ and K_2_. Under low-field and diffusion-limited conditions, apex separation grows linearly with the drift length L. Longitudinal diffusion, however, broadens each peak width but only as L^1/2^. Combining these two effects, the resolution is expected to be a function of the square root of n (SI, Experimental Section).

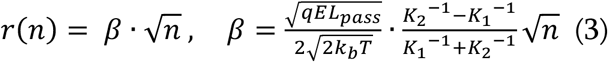

Where q is the charge of isomer, kb is the Boltzmann’s constant, T is the temperature, and E is the electric field. Figure 5A illustrates the progressive separation of the triply charged H3_[3-15]_K9bu/ibu pair as n increases from 1 to 60. Each species (individual isomers and their mixture) is analyzed in an independent multipass experiment, and within each experiment drift times were recorded sequentially across pass numbers. Short-term repeatability can be assessed by comparing the drift-time apex of each isomer measured individually with its corresponding apex observed in the mixture. The relative drift-time deviation (Δdt/<dt>) for this pair is 0.1% (0.1 ms at 100 ms drift time), indicating stable cIM performance under identical conditions. Regarding, longer-term repeatability is evaluated for H3_[3–25]_K9K18 doubly modified peptides measured 444 or 456 days apart (Figure S11). These experiments yield relative drift time deviations of 1.18 % for single pass experiments and 0.16 % for 40 passes, respectively, thus demonstrating sufficient long-term stability for reliable referencing of drift time values against reference spectra.

**Figure 5.**
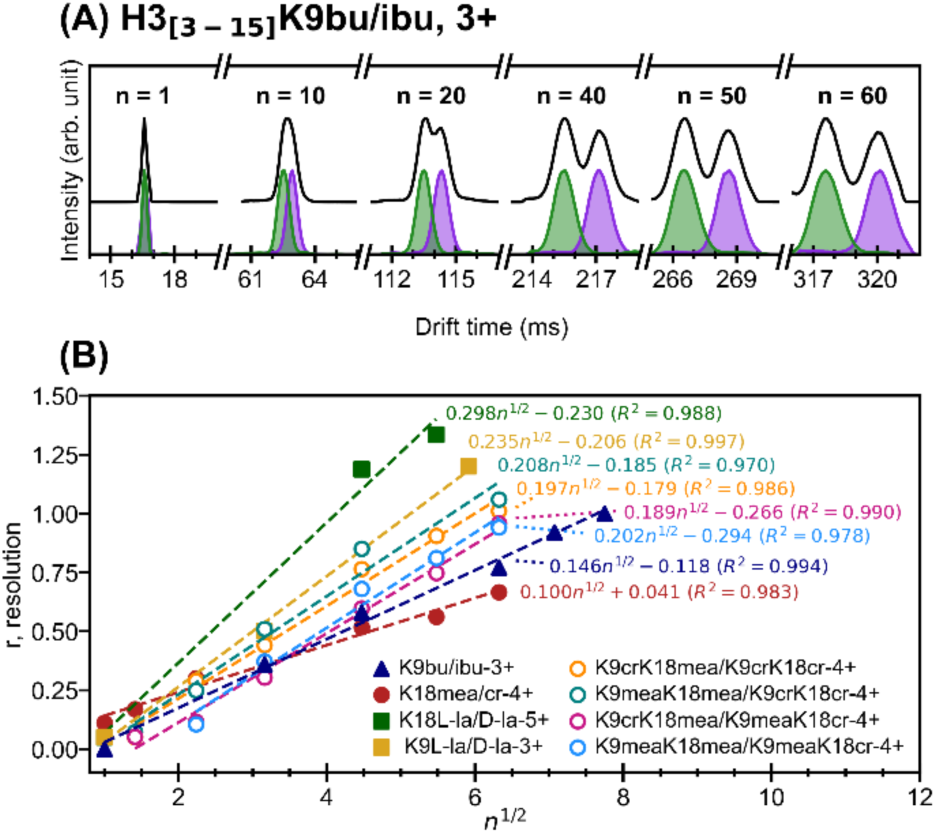
Resolution scaling in multpass cIM. (A) Progressive cIM–MS separation of triply charged H3_[3-15]_K9bu (violet) and K9ibu (green) as the path length increases from 1 to 60 passes. (B) Experimental resolution (r) plotted as a function of the square root of the number of passes (n^1/2^). Filled and open symbols represent singly and doubly modified species, respectively. The dotted lines are least-squares linear fits to each dataset.

Figure 5B plots r as a function of n^1/2^ for several other isomeric pairs where drift time distributions were recorded for different n values. These datasets follow a linear dependency with R_2_ > 0.97, confirming experimentally the dependency of r on the square root of n. In practice, once a measurable apex split is observed, the equation (3) provides a simple rule-of-thumb where the number of passes required for baseline separation can be estimated. Importantly, the linear n^1/2^ relationship reveals that the ions remain conformationally static over the experimental timescale, with negligible contributions from interconverting or alternative conformers to each mobility peak and no detectable structural perturbation during the multipass separation.

## CONCLUSION

Histone proteins undergo diverse PTMs, some of them leading to isomeric acyl-lysine marks with important roles in human biomedicine.^4, 20^ Separation and identification of isomeric PTMs using biochemical and bioanalytical techniques has remained a challenging task, thus development of novel mass spectrometric tools addressing these challenges has recently gained a widespread interest from the biochemical and chemical community. Here we demonstrate that cyclic IMS achieves baseline or near-baseline resolution for every isomeric crotonyl/methacryl, butyryl/isobutyryl and L-/D-lactyl pair of histone H3 peptides examined, with exception of long H3_[3-25]_K18bu/ibu isomers, which remain unseparated after 60-70 passes. TIMS achieves half-height separation for L-/D-lactyl- and crotonyl-/methacryl-modified H3_[3-15]_ peptides, and the H3_[3-25]_K18la peptides. Across all separated isomeric PTM species, average CCS/drift time differences are ∼0.5-1% for L/D-lactyl, ∼0.8% for butyryl/isobutyryl, and ∼0.4% for crotonyl/methacryl for H3_[3-15]_ peptides. Corresponding values of ∼1.0% (L/D-lactyl) and ∼0.4-0.8% (crotonyl/methacryl) are measured for H3_[3-25]_K18 peptides. Systematic trends are observed: branched acyl groups (methacryl, isobutyryl) consistently produce more compact conformers than their linear counterparts, and the CCS difference is amplified in doubly modified peptides. Separation order of L- and D-lactyl isomers in IMS spectra is charge state dependent and resolution in cIM follows the expected n^1/2^ scaling with number of cIM passes. The IMS separations reported here occur within ∼300-1300 ms in high-resolution IMS scans, these durations that remain compatible with typical LC peak widths. This has been demonstrated experimentally in targeted LC–TIMS–MS analyses of isomeric N-methylhistidine peptides, where 1.3 second TIMS ramps enabled confident isomer identification in a complex HeLa digest.^51^ Thus, the present separations are transferable to targeted LC–IMS–MS/MS workflows and for extending this approach to endogenous modified histone proteins and their proteolytically produced peptides.

Unambiguous identification of isomeric PTMs with IMS benefits from the comparison to reference spectra. Targeted ozone-induced dissociation can provide complementary structural evidence for the unsaturated acyl modification.^9^ Further improvement in (isomeric) PTM assignments will likely rely on predictive computational models. Fine-tuning existing machine-learning based CCS models using high-quality IMS datasets, including cIM, TIMS, drift-tube IMS measurements of wider range of acyl-PTM isomers, will enable prediction and validation below the sub-percent CCS differences observed here. This approach has an exceptional potential for de-novo assignment of PTM isomers across diverse proteins.

## ASSOCIATED CONTENT

### Supporting Information

Detailed experimental procedures (peptide synthesis, sample preparation, and infusion conditions) and instrument settings for TIMS and cyclic IMS (scan rates, pass numbers/path lengths, and CCS calibration) are provided. Additional IMS–MS data for all isomer pairs and charge states (Figures S1–S11). The n^1/2^ dependence of peak-to-peak resolution. Table S1 summarizes sequences and modification sites.

## AUTHOR INFORMATION

### Author Contributions

The manuscript was written through contributions of all authors. All authors have given approval to the final version of the manuscript.

### Notes

The authors declare the following competing financial interest: DACS and JIL are employees of Waters Corporation, which manufactures the cyclic ion mobility instrument used in this study. The other authors declare no competing financial interest.

## ACKNOWLEDGMENT

This research was supported by the Independent Research Fund Denmark (013500114B to O.N.J.), the Lundbeck Foundation (R344-2020-1051 to J.M.) and the Novo Nordisk Foundation (NNF19OC0058779 to J.M., and NNF20OC0061575 to O.N.J.).

## TABLE OF CONTENTS GRAPHIC

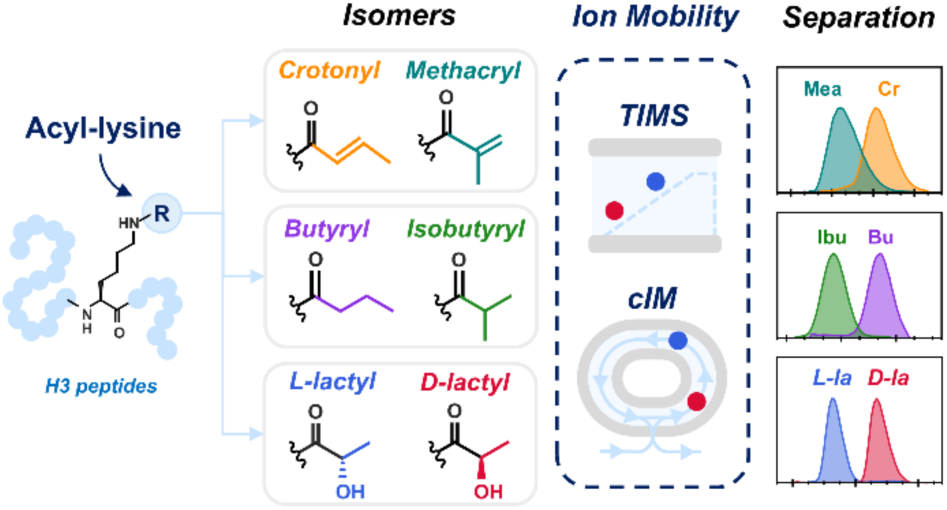

