## Supplementary Information for "Ion Mobility Separation of Isomeric Acyl-lysine Marks in Peptides"

§Waters Corporation, Stamford Avenue, Altrincham Road, Wilmslow SK9 4AX, Cheshire, U.K.

---

TABLE OF CONTENTS.

**Experimental Section**
- Page S2-S3: Peptide synthesis, acyl-lysine installation, sample preparation, and IMS-MS instrument settings for TIMS and cyclic IMS.
- Page S4: Peptide Overview. Table S1 summarizing sequences, modification sites, and theoretical masses of all synthetic histone H3 peptides.
- Page S5-S10: Ion mobility instrumentation and resolution metrics. Operating principles and experimental settings for TIMS and cyclic IMS, definitions of IMS parameters, resolution metrics, and derivation of resolution scaling with the number of cyclic IMS passes.

**Supplementary Figures**
- Page S11-S21: Supplementary Figures: Figures S1–S11 showing additional TIMS-MS and cIM-MS spectra (Figures S1-10), and long-term drift-time stability (Figure S11).

EXPERIMENTAL SECTION

**1. Peptides synthesis.**

Histone H3 peptides were synthesized utilizing automated Fmoc-SPPS chemistry on Rink amide resin (0.78 mmol/g loading capacity) on a PurePep® Chorus Synthesizer. Coupling of the amino acids (3.0 eq.) was carried out at 75 °C for 20 min upon activation with HATU (3.0 eq.) and DIPEA (4.5 eq) in DMF. Arg was coupled twice for 15 min. Fmoc deprotection was achieved with a solution of 20 % piperidine in DMF for 7 min at 50 °C. Alloc-protected lysine residues were incorporated at K9 and K18 positions. The last amino acid in the sequence was coupled as Boc-Thr. Upon completion of the sequence, the lysine residues were selectively modified with the different acyl groups on-resin. Initially, the resin containing fully protected H3 peptides was swollen in DCM under nitrogen flow for 10 min at room temperature. The on-resin peptide proceeded to orthogonal removal of the alloc-protecting group by treatment with $PhSiH_3$ (20 eq.) and $Pd(PPh_3)_4$ (1.0 eq.) to the $N_2$-bubbling resin in DCM for 1 h at room temperature. Extensive washing of the resin with DCM, DMF and sodium diethyldithiocarbamate (0.5 % in DMF, w/v) was followed by a Kaiser test to monitor deprotection. Acylation reactions were then carried out on H3K9 and H3K18 at different conditions.

*Crotonyllysine/Methacryllysine peptides:* For the single-modified peptides, triethylamine (TEA) (1.5 eq.) in DCM was added to the resin and agitated for 10 min at room temperature. Subsequently, *trans*-crotonyl chloride (3.0 eq) or methacroloyl chloride (3.0 eq.) was added to the resin and the reaction was allowed to proceed for 2 h and monitored by Kaiser test. For doublemodified peptides, the resin was first elongated to position 18 (including residues 18–25), followed by orthogonal alloc deprotection and selective lysine acylation as described above. Elongation was then continued, and the same procedure was applied at position 9.

*Butyryl/Isobytyryllysine peptides:* For the installation of butyryl or isobutyryl modifications, the resin was first treated with TEA (1.5 eq.) in DCM and agitated for 10 min at room temperature. Subsequently, butyryl chloride (3.0 eq.) or isobutyryl chloride (3.0 eq.) was added to the resin. The acylation was allowed to proceed for 1 hour under continuous agitation, and reaction completion was monitored by the Kaiser test.

*L-/D-Lactyllysine peptides:* L-(+)-lactic acid or D-(–)-lactic acid was dissolved in DCM and activated with HATU and HOBt (2.9 and 3.0 eq., respectively) in the presence of DIPEA (5.0 eq.) using a DCM:DMF (2:1) solvent mixture. The resulting solution was added to the resin and the coupling reaction was carried out under agitation for 1 hour, with progress monitored by the Kaiser test. This procedure was repeated for each acylating reagent.

After completion of the acylations, the final peptides were washed with DCM, dried over diethyl ether, and then proceeded to cleave from resin using 95% (v/v) TFA, 2.5% (v/v) TIPS, and 2.5% (v/v) MQ-water for 4 h. The crude peptides were precipitated with cold diethyl ether (–20 °C) and pelleted via centrifugation. Finally, the synthetic modified histone peptides were purified by RP-HPLC, and their purity was assessed by MALDI-TOF MS and analytical RP-HPLC.

**Table S1.** Overview of all synthetic peptides

| Peptides | Sequence[(a)] | Theoritical monoisotopic neutral mass (Da) |
|---|---|---|
| $H3_{[3-15]}$K9cr | TKQTARK*STGGKA | 1399.78 |
| $H3_{[3-25]}$K18cr | TKQTARKSTGGKAPRK*QLATKAA | 2464.43 |
| $H3_{[3-15]}$K9mea | TKQTARK*STGGKA | 1399.78 |
| H3K18mea | TKQTARKSTGGKAPRK*QLATKAA | 2464.43 |
| $H3_{[3-25]}$K9meaK18mea | TKQTARK*STGGKAPRK*QLATKAA | 2532.45 |
| $H3_{[3-25]}$K9meaK18cr | TKQTARK*STGGKAPRK*QLATKAA | 2532.45 |
| $H3_{[3-25]}$K9crK18mea | TKQTARK*STGGKAPRK*QLATKAA | 2532.45 |
| $H3_{[3-25]}$K9crK18cr | TKQTARK*STGGKAPRK*QLATKAA | 2532.45 |
| $H3_{[3-15]}$K9L-la | TKQTARK*STGGKA | 1403.78 |
| $H3_{[3-15]}$K9D-la | TKQTARK*STGGKA | 1403.78 |
| $H3_{[3-25]}$K18L-la | TKQTARKSTGGKAPRK*QLATKAA | 2468.43 |
| $H3_{[3-25]}$K18D-la | TKQTARKSTGGKAPRK*QLATKAA | 2468.43 |
| $H3_{[3-15]}$K9ibu | TKQTARK*STGGKA | 1401.80 |
| $H3_{[3-15]}$K9bu | TKQTARK*STGGKA | 1401.80 |
| $H3_{[3-25]}$K18ibu | TKQTARKSTGGKAPRK*QLATKAA | 2466.45 |
| $H3_{[3-25]}$K18bu | TKQTARKSTGGKAPRK*QLATKAA | 2466.45 |

**(a)** an asterisk marks the modified residue.

### 2. Ion mobility instrumentation and resolution metrics

| Symbol | Meaning | Unit |
|---|---|---|
| $CCS$ | Collision Cross Section | $m^2$ |
| $D$ | Diffusion coefficient | $m^2 \cdot s^{-1}$ |
| $E$ | Electric field | $V \cdot m^{-1}$ |
| $FWHM$ | Full width at half-maximum | s |
| $K$ | Ion mobility coefficient | $m^2 \cdot V^{-1} \cdot s^{-1}$ |
| $L$ | Total drift length | m |
| $L_{pass}$ | Path length per pass (~1m) | m |
| $N$ | Number density of the drift gas | $m^{-3}$ |
| $T$ | Temperature | K |
| $e$ | Elementary charge | C |
| $k_n$ | Boltzmann's constant | $J \cdot K^{-1}$ |
| $n$ | Number of passes | |
| $q$ | Ion charge | C |
| $r$ | Resolution | |
| $R_p$ | Resolving power | |
| $Sr$ | TIMS scan rate | $V \cdot s^{-1}$ |
| $\sigma$ | Temporal diffusion | s |
| $\sigma_s$ | Spatial diffusion | m |
| $t_d$ | Drift time | s |

#### 2.1 Trapped IMS-MS

*Separation principle:* Ions are entrained in a buffer gas flow and radially confined by application of a radiofrequency field along the TIMS tunnel. Under the influence of a moderate electric field (E) oriented in the opposite direction of the propagation to the ions, the ions with different mobilities (K) are trapped at different equilibrium positions where the forward drag force is counterbalanced by the deceleration force imposed by E. Ions with higher mobility are trapped closer to the entrance, while ions with lower K are trapped further downstream. Separation is achieved by ramping the electric field down in a controlled ramp (scan rate, Sr), releasing ions in order of increasing mobility.

*Settings:* Trapped IMS experiments were performed on a timsTOF Pro 2 hybrid Q-TOF instrument (Bruker Daltonics). All experiments were performed in positive-ion mode Protonated synthetized peptides were directly infused via a CaptiveSpray nanoESI source (Bruker Daltonics) at a final concentration of 1-3 μM, in water/methanol/formic acid (49:50:1) solvent. The settings of the ESI source were capillary voltage = 1.1 kV, Dry gas flow rate = 3 L/min and dry gas temperature set up at 180°C.

*Calibration:* Reduced inverse mobility coefficient ($K_0^{-1}$) and mass-to-charge ratios (*m/z*) were calibrated using ESI-L low-concentration tune mix (Agilent). Collision cross sections (CCS) using $N_2$ as buffer gas were calculated from calibrated mobilities via the Mason–Schamp equation, based on literature reference values for protonated fluorinated phosphazine ions within the tune mix: *m/z* 622.03 ($K_0^{-1}$ = 0.992 V·s·cm$^{-2}$, CCS = 203.0 Å$^2$), *m/z* 922.01 (1.198 V·s·cm$^{-2}$, 243.6 Å$^2$):

$$CCS = \frac{3}{16}\sqrt{\frac{2\pi}{\mu k_b T}}\frac{ze}{NK}$$

where μ is the reduced mass of the colliding partners. Since K varies with the pressure (P) and the temperature (T), it is commonly corrected to the reduced mobility $K_0$, normalized to standard conditions ($T_0$ and $P_0$, respectively):

$$K_0 = K\frac{P \cdot T_0}{P_0 \cdot T}$$

*Resolving power improvement.* Trapped IMS settings were optimized for high resolving power, as described previously.[1, 2] Briefly, the in-tunnel TIMS pressure was increased to 3.4 mbar (vs. 2.6 mbar standard), and the TIMS scan rate (Sr) was decreased by employing a 1.3 s TIMS ramp and applying focused ion-mobility windows around regions of interest. The accumulation time was set at 40 ms prior to TIMS analysis.

#### 2.2 Cyclic IM-MS

*Separation principle.* The cIM analyzer operates on the same principle as travelling wave IMS (TWIMS) but with a closed-loop drift path. Gas-phase ions are guided through a closed-loop stacked ring ion guide of 0.98 m path length. The ions move axially along the drift tube in response a sequence of voltage pulses (the “traveling waves”) through a drift gas. Ions with different mobilities travel with different effective velocities ($v_d$) relative to the travelling wave, and thus separate temporally. A superimposed RF voltage provides radial confinement, ensuring ion transmission. Ions exit the cIM analyzer in the same order as their drift time, which reflects their gas-phase mobility.

*Settings:* Cyclic IMS-MS experiments were performed on an ESI-Q-cIM-ECD-TOF instrument (Waters). IMS settings used included a traveling wave (TW) velocity of 375 m/s and a TW static height of 22 V. The drift gas is $N_2$ at a pressure of 1.8 mbar Synthetized peptides were directly infused at a final concentration of 1-3 µM, in water/methanol/formic acid (49:50:1) solvent, using an ESI voltage of 1.2 kV.

*Resolving power improvement.* The circular geometry of the cIM analyzer enables an adaptive resolving power that can be increased by increasing the number of passes. Specific ions can be isolated based on their arrival times for additional passes using the top-and-tail method, which prevents high-mobility ions from catching up with low-mobility ions. In a cIM analyzer, the resolving power scales with the square root of the number of passes.[3]

#### 3. Resolving power and resolution

The resolving power, Rp, was determined from the apex position of a peak in an IMS spectrum and from its full width at half maximum (FWHM) using Compass DataAnalysis software (Bruker Daltonics, v6.1) for TIMS-MS data and MaxLynx software (Waters, v4.2) for cIM-MS data.

$$R_{p=}\frac{x_i}{FWHM_i}$$

Where $x_i$ is the CCS, $K_0^{-1}$ or $t_d$ value at the apex of peak i. The resolution, r, also referred as peak-to-peak resolution, was calculated assuming Gaussian peak shapes:

$$r\ = 1.178 \cdot \frac{|x_2 - x_1|}{FWHM_1 + FWHM_2}$$

The resolution quantifies the separation between species, and a baseline separation is defined with a r value greater than ~1.1, half-height separation corresponds to r ~ 0.7.

#### 4. Resolution as a function of the number of passe (multipass cIM).

We consider the low-field IMS regime, where the ion mobility K is independent of the electric field E. In addition, we consider all operating conditions (gas temperature and pressure, travelling-wave amplitude and speed) remain constant across passes. IMS peaks are assumed to be Gaussian, and longitudinal diffusion is the dominant source of temporal dispersion. For an ion of mobility K, the drift velocity ($v_d$) is:

$$v_d = KE$$

The drift time for the total drift length (n passes) is:

$$t_d = \frac{L}{KE} = \frac{nL_{pass}}{KE}$$

The drift time difference between ions 1 and 2 is:

$$\Delta t_d = t_{d,2} - t_{d,1} = \frac{nL_{pass}}{E}\left(\frac{1}{K_2} - \frac{1}{K_1}\right)$$

The temporal diffusion, $\sigma$, of an ion i is due to random collision causing ion clouds to spread.

Longitudinal diffusion in one dimension produces a spatial variance (Ficks' law):

$$\sigma_{s,i}{}^2 = 2Dt_{d,i}$$

The corresponding temporal dispersion, $\sigma$, is obtained by dividing the $\sigma_s$ by the drift velocity, giving:

$$\sigma_i = \sqrt{\frac{2D_i nL_{pass}}{(K_i E)^3}}$$

In IMS, the resolution is defined by analogy to chromatography:

$$r = \frac{t_{d,2} - t_{d,1}}{\frac{w_1 + w_2}{2}} = \frac{2 \cdot \Delta t_d}{w_1 + w_2}$$

Where $w_i$ is the width at the base of a peak i. By convention and assuming gaussian shape peaks $w$ is egal to $4\sigma$ .

The resolution can be expressed as a function of the number of passes for two isomers 1 and 2:

$$r = \frac{\sqrt{E \cdot L_{pass}}}{2\sqrt{2}} \cdot \frac{K_2^{-1} - K_1^{-1}}{\sqrt{D_1} \cdot K_1^{-\frac{3}{2}} + \sqrt{D_2} \cdot K_2^{-\frac{3}{2}}} \sqrt{n}$$

The Einstein–Smoluchowski relation gives the relation between diffusion and mobility:

$$D_i = \frac{k_B T}{q} K_i$$

Finally, the resolution as a function of n is thus expressed as:

$$\boxed{r = \frac{\sqrt{qEL_{pass}}}{2\sqrt{2k_b T}} \cdot \frac{K_2{}^{-1} - K_1{}^{-1}}{K_1{}^{-1} + K_2{}^{-1}} \sqrt{n}}$$

In Figure 5, the experimental r values were fitted using a linear function $r(n)=an^{1/2} + b$ rather than forcing b=0, to consider any n-independent contributions to the peak width (e.g., injection dispersion, gate jitter, …) that shift the intercept from the ideal diffusion-limited case.

### SUPPLEMENTARY FIGURES

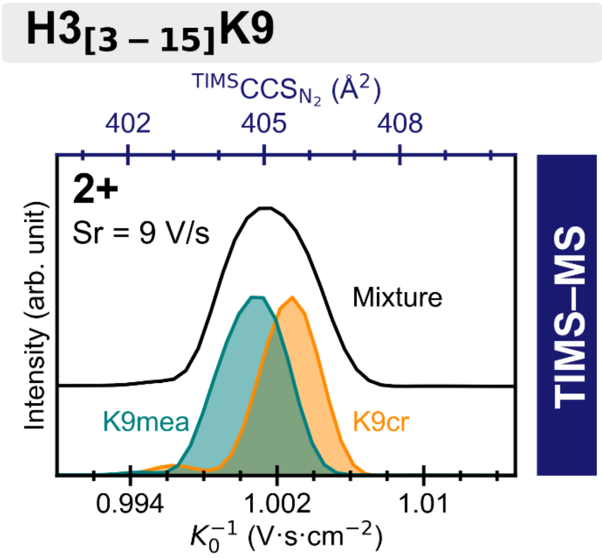


**Figure S1.** TIMS–MS spectra of $H3_{[3\text{-}15]}$K9mea (teal) and $H3_{[3\text{-}15]}$K9cr (orange) at charge state 2+ (Sr = 9 $V{\cdot}s^{-1}$). The mixture trace is shown in black.

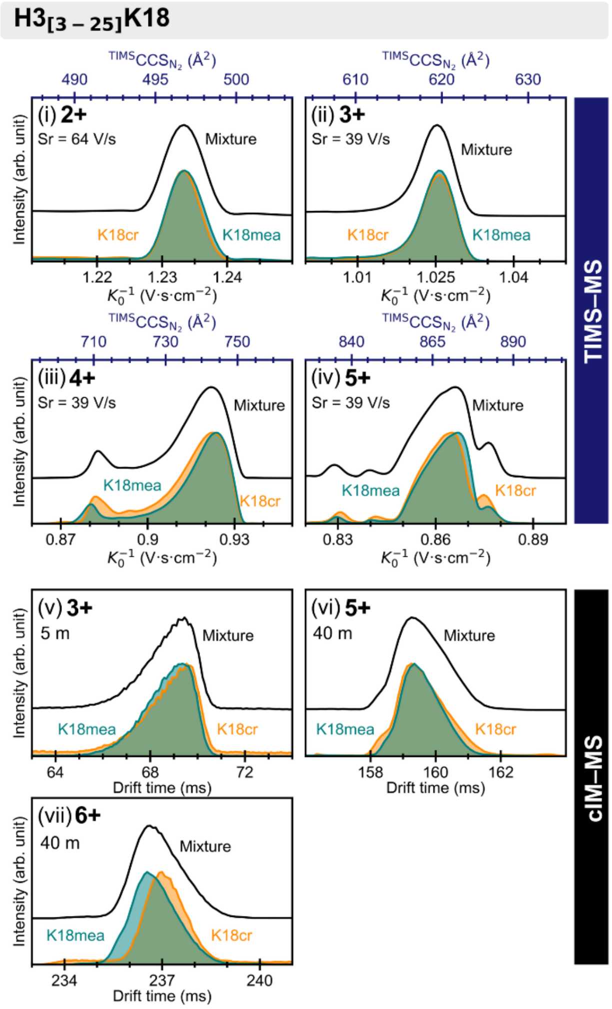


**Figure S2**. IMS–MS spectra of H3[3-25]K18mea (teal) and H3[3-25]K18cr (orange). (i–iii) TIMS–MS spectra of charge state 2+ (Sr = 64 V·s$^{-1}$), and charge states 3+–5+ (Sr = 39 V·s$^{-1}$). (iv–vii) cIM drift time distributions of charge states 4+, 5+, and 6+ recorded after 5, 40, and 40 passes, respectively. The mixture traces are shown in black.

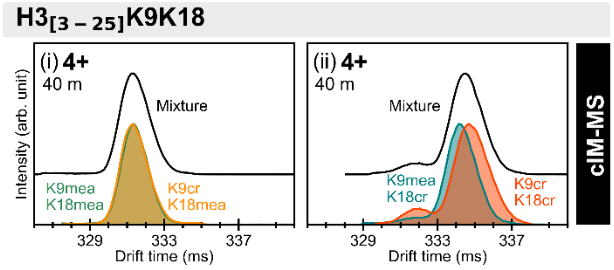


**Figure S3.** cIM–MS drift-time distributions for doubly modified H3[3-25]K9K18 peptides bearing crotonyl and methacryl groups. (i) K9meaK18cr vs. K9crK18mea; (ii) K9meaK18cr vs. K9crK18cr for charge state 4+ after 40 passes. The mixture traces are shown in black.

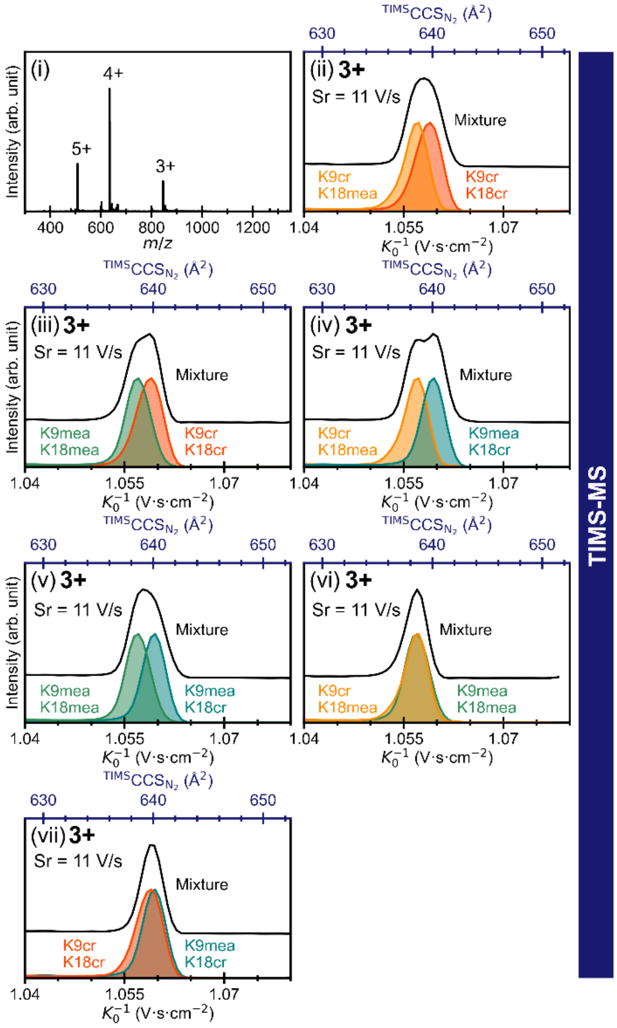


**Figure S4**. TIMS–MS distributions of doubly modified H3[3-25]K9K18 peptides bearing crotonyl and methacryl groups. (i) Mass spectrum; (ii–vi) TIMS–MS distributions of charge states 3+ recorded at Sr = 11 V·s$^{-1}$. The mixture traces are shown in black.

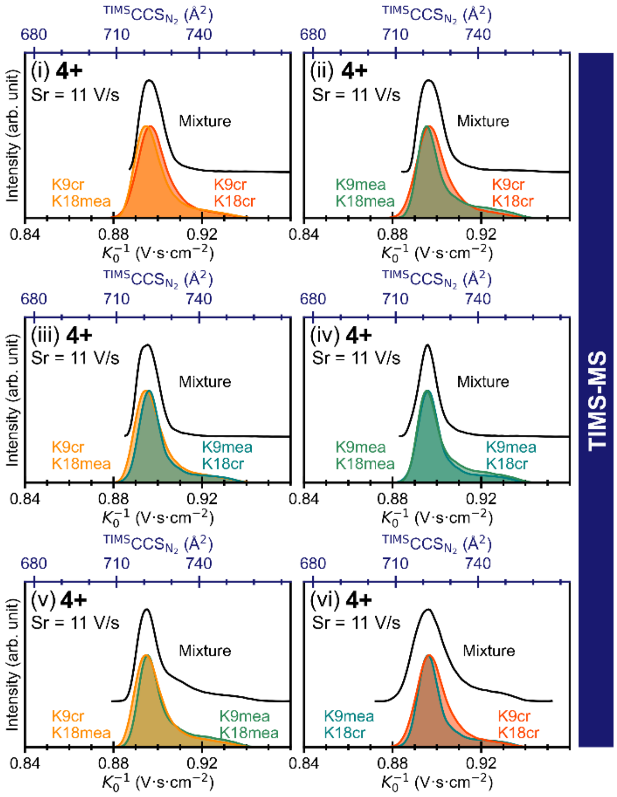


**Figure S5**. TIMS–MS distributions of doubly modified H3[3-25]K9K18 peptides bearing crotonyl and methacryl groups (i–vi) TIMS–MS distributions of charge states 4+ recorded at Sr = 11 V·s$^{-1}$. The mixture traces are shown in black.

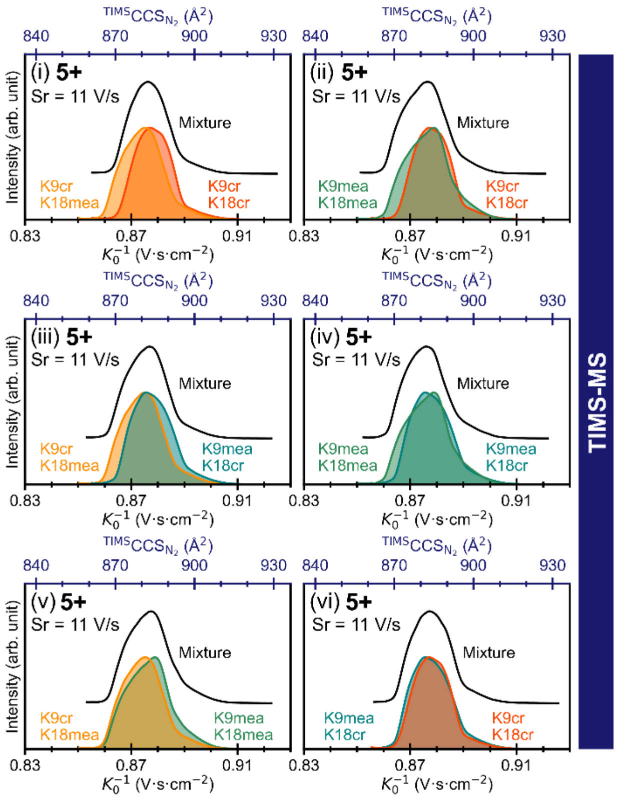


**Figure S6**. TIMS–MS distributions of doubly modified H3[3-25]K9K18 peptides bearing crotonyl and methacryl groups. (i–vi) TIMS–MS distributions of charge states 5+ recorded at Sr = 11 V·s$^{-1}$. The mixture traces are shown in black.

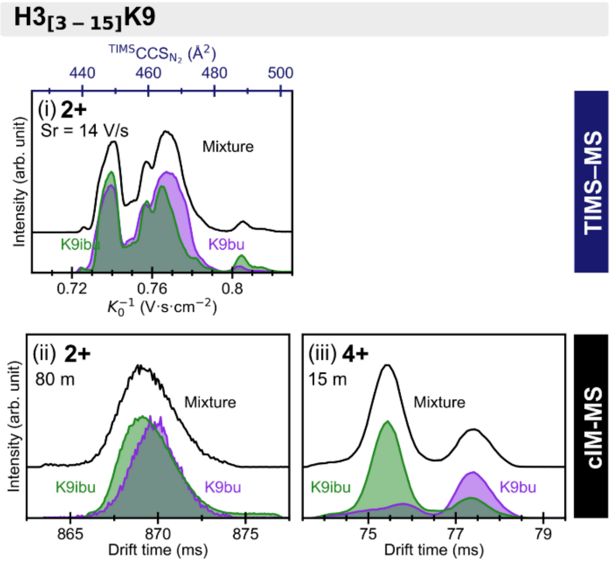


**Figure S7.** IMS-MS distributions of H3[3-15]K9bu (violet) and H3[3-15]K9ibu (green). (i-ii) TIMS-MS spectra of charge states 2+ and 3+ (Sr = 14 V·s$^{-1}$). (iii) and (iv) cIM distributions of charge states 2+ and 4+ after 80 and 15 passes, respectively. The mixture traces are shown in black.

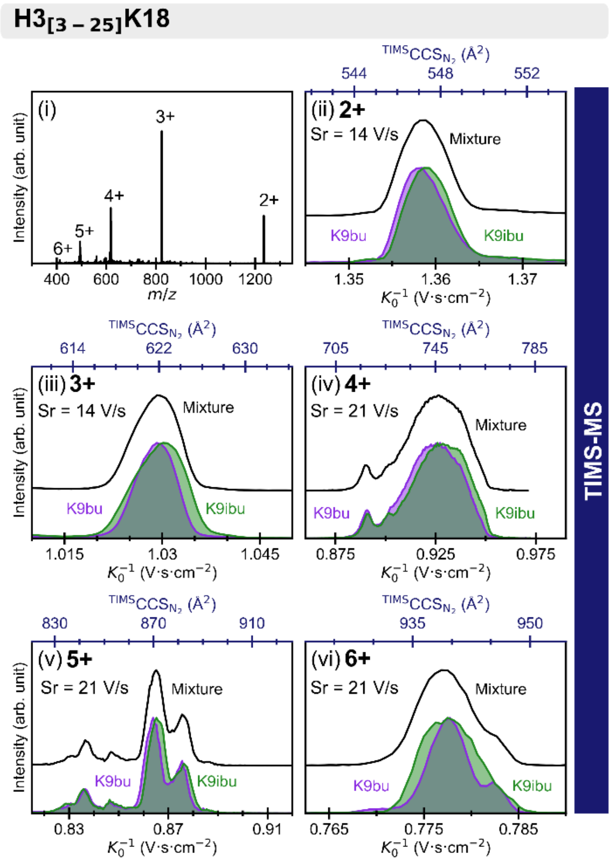


**Figure S8**. TIMS–MS distributions of H3[3-25] K18bu (violet) and K18ibu (green). (i) Mass spectrum; (ii–iii) TIMS–MS distributions of charge states 2+ and 3+ recorded at Sr = 14 V·s$^{-1}$. (iv–vi) TIMS–MS distributions of charge states 4+, 5+, and 6+ recorded at Sr = 21 V·s$^{-1}$. The mixture traces are shown in black.

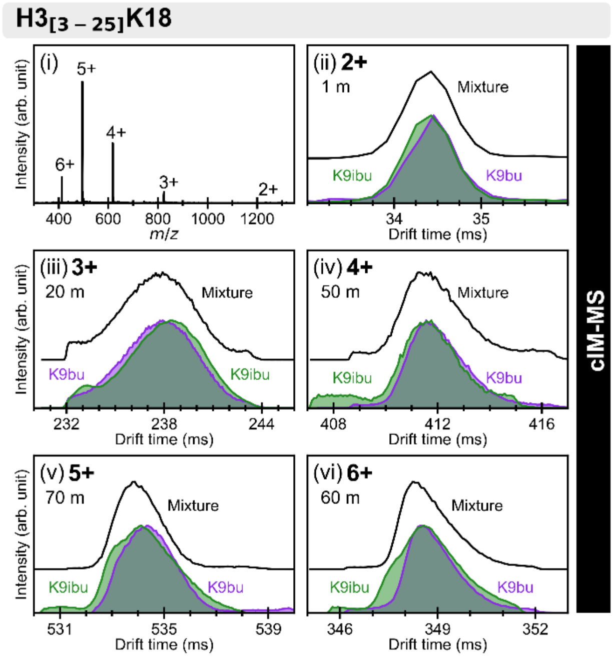


**Figure S9**. cIM–MS drift time distributions of H3[3-25]K18bu (violet) and H3[3-25]K18ibu (green). (i) Mass spectrum; (ii–vi) cIM distribution of charge states 2+, 3+, 4+, 5+ and 4+ after 1, 20, 50, 70 and 60 passes, respectively. The mixture traces are shown in black.

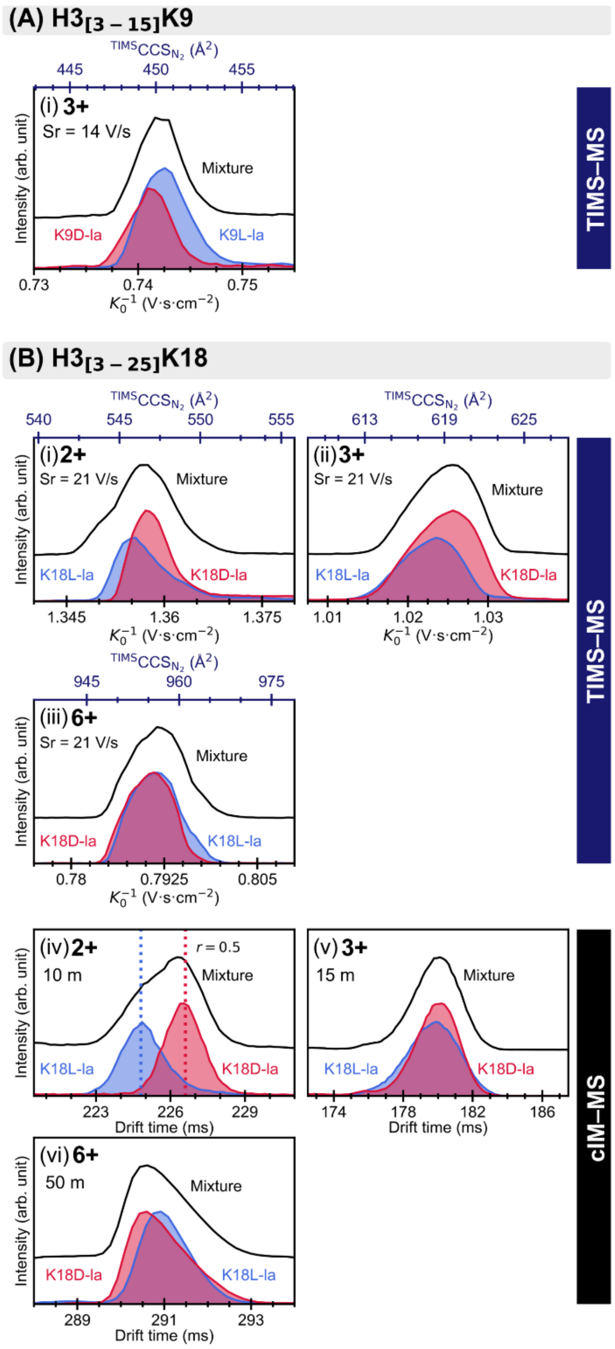


**Figure S10.** IMS-MS distributions for lactyl isomers at K9 ($H3_{[3\text{-}15]}$) and K18 ($H3_{[3\text{-}25]}$). (A) $H3_{[3\text{-}15]}$K9L-la/K9D-la: TIMS distributions of charge state 3+ for $H3_{[3\text{-}15]}$K9L-la/D-la (Sr = 14 V·s$^{-1}$). (B) $H3_{[3\text{-}25]}$K18L-la/K18D-la: (i-iii) TIMS distributions of charge states 2+, 3+, and 6+ (Sr = 21 V·s$^{-1}$);

(iv-vi) cIM distributions of charge states 2+, 3+, and 6+ after 10, 15, and 50 passes, respectively. KL-la in blue, KD-la in red and mixtures in black.

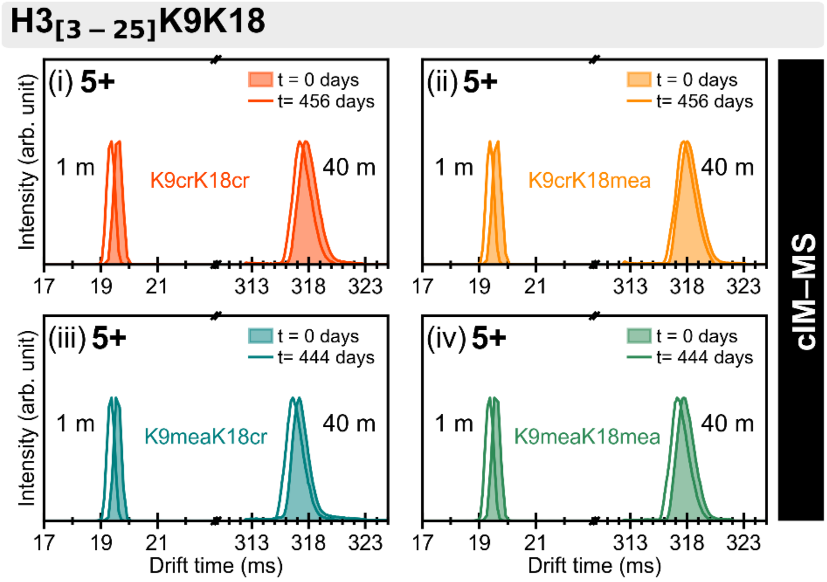


**Figure S11.** Long-term cIM drift-time stability for doubly modified $H3_{[3-25]}$K9K18 peptides at charge state 5+. Drift-time distributions of $H3_{[3-25]}$K9crK18cr (i), $H3_{[3-25]}$K9crK18mea (ii), $H3_{[3-25]}$K9meaK18cr (iii), and $H3_{[3-25]}$K9meaK18mea (iv)c on two measurements separated by 444 or 456 days. The relative drift-time deviations (Δdt/<dt>) between day 0 and day 444/456 measurements were estimated to 1.18% for single-pass data (corresponding to 0.23 ms at a 20 ms drift time) and 0.16% for 40-pass data (0.5 ms at a 320 ms drift time).
